# Enteropathogenic *Providencia alcalifaciens*: A subgroup of *P. alcalifaciens* that causes diarrhea

**DOI:** 10.1101/2024.04.26.591277

**Authors:** Dieter Bulach, Glen P. Carter, M. John Albert

## Abstract

Even though *Providencia alcalifaciens* is considered as a normal flora of the large intestine, there are reports of it causing diarrhea. In a previous study, a strain, 2939/90 obtained as a pure stool culture from a dead diarrheal patient was shown to cause invasion and actin condensation in mammalian cells, and diarrhea in a rabbit model. In a subsequent study, four Tn*phoA* mutants of 2939/90 produced negligible invasion and actin condensation in mammalian cells. In the present study, the parent strain was sequenced by short-read and long-read sequencing, and the mutants by short read sequencing. In all four mutants, a Tn*phoA* insertion was detected in the type three secretion system (T3SS) locus present on the largest of four plasmids (p2939_90_1) and not in a seemingly independent, functional T3SS locus on the chromosome. A survey of 52 genomes of *P. alcalifaciens* available in the public database identified the chromosomal T3SS locus in all strains, including both *P. alcalifaciens* genomic clades that we have classified as (group A) and (group B); a highly related gene layout and gene synteny flanking the locus suggested that these chromosomal loci are orthologous. There is a low sequence similarity between the chromosomal and plasmid-borne T3SS; a survey of plasmid T3SS showed its presence in only 21 of 52 genomes and mostly in group A genomes. Group A included several isolates from an outbreak of haemorrhagic diarrhea in dogs. Using prediction software (EffectiveDB), we detected several known and unknown effectors flanking the plasmid T3SS locus. The observation that Tn*phoA* insertion only in the plasmid T3SS locus affected the invasion phenotype suggested that this locus is critical for causation of diarrhea. This leads us to conclude that a subgroup of *P. alcalifaciens* that possesses this plasmid-borne T3SS locus (in the case of strain, 2939/90) can cause diarrheal disease. We name this subgroup as enteropathogenic *P. alcalifaciens* (EPA). EPA should be included in future studies of etiology of diarrhea. A unique sequence that may be present in the T3SS locus in the plasmid may be investigated as a marker in a simple molecular test for diagnosis of EPA.

## Introduction

*Providencia alcalifaciens* is a species in the *Providencia* genus of the family *Enterobacteriaceae* (Janda and Abbott, 2006). Although *P. alcalifaciens* is considered a part of the normal flora of the feces in many mammals including humans, there is evidence to suggest that it can also cause diarrhea. It has been implicated in foodborne outbreaks of diarrhea (Murata et al, 2001; Shah et al, 2015) and travellers’ diarrhea (Haynes and Hawkey, 1989). In a case-control study of children’s diarrhea, the organism was isolated at a significantly higher rate from children with diarrhea than from matched control children (Albert et al, 1998). It was reported to cause foodborne haemorrhagic diarrhea in dogs (Jorgensen et al, 2021). Pathogenicity studies with diarrheal isolates suggested that *P. alcalifaciens* can invade cultured mammalian cells such as HEp-2 cells with actin condensation, and cause diarrhea in reversible ileal tie adult rabbit diarrhea (RITARD) model with the invasion of the intestinal mucosa (Albert et al, 1992). There were two modes of invasion of epithelial cells: one by endocytosis and the other through intercellular tight junction (Mathan et al, 1993). Tissue culture invasion was inhibited by an agent that prevented microfilament formation (Albert et al, 1995). Further studies confirmed that many diarrheal isolates of *P. alcalifaciens* were invasive for mammalian cells, but some were not (Beatriz et al, 1996; Janda et al, 1998). To define the basis of invasion, Tn*PhoA* mutagenesis of the diarrheal strain, *P. alcalifaciens* 2939/90, was used to demonstrate cell invasion and actin condensation. The Tn*PhoA* mutants exhibited negligible invasion and actin condensation in HEp-2 cell assays (Rahman et al, 2002). In the current study, the parent strain, 2939/90 and four Tn*PhoA* insertion mutants were sequenced to determine the insertion sites of Tn*PhoA* and elucidate the genetic basis of virulence. Evaluation of the distribution of genetic determinants contributing to cell invasion in strain 2939/90 and across the *P. alcalifaciens* species leads us to hypothesize that a specific lineage of *P. alcalifaciens* is diarrheagenic. We refer to this subgroup that causes diarrhea as enteropathogenic *P. alcalifaciens*.

## Materials & Methods

### Isolates

We studied the parent wildtype strain of *P. alcalifaciens* 2939/90 that was isolated from a child with diarrhea who was dead on arrival at a hospital in Dhaka, Bangladesh. This strain had an invasive phenotype for the intestine in an animal model of diarrhea and in an *in vitro* HEp-2 cell assay (Albert et al., 1992). In addition, four Tn*PhoA* mutants of *P. alcalifaciens* 2939/90 that had negligible invasion in HEp-2 cells (Rahman et al., 2002) were studied. The details of the isolates are shown in Table 1.

**Table 1.** *P. alcalifaciens* strain 2939/90 and its Tn*PhoA* mutants sequenced in this study.

| Strain/Mutant | Genome Sequence | Invasive phenotype<sup>^</sup> | Reference |
| --- | --- | --- | --- |
| 2939/90 | Chromosome & four Plasmids | Yes | Albert et al., 1992 |
| M-23_Tn*PhoA* | Contigs | Negligible | Rahman et al., 2002 |
| M-47_Tn*PhoA* | Contigs | Negligible | Rahman et al., 2002 |
| M-63_Tn*PhoA* | Contigs | Negligible | Rahman et al., 2002 |
| M-78_Tn*PhoA* | Contigs | Negligible | Rahman et al., 2002 |

<sup>^</sup>Parent tested in animal model and Hep-2 cells, and mutants tested in Hep-2 cells

**Table 2.** Core genome, pairwise SNP distance between parental strain (2939/90) and Tn*PhoA* mutants.

| Strain/Mutant | 2939/90 | M-23* | M-47* | M-63* | M-78* |
| --- | --- | --- | --- | --- | --- |
| 2939/90 | 0 | 1 | 2 | 1 | 1 |
| M-23 | 1 | 0 | 1 | 0 | 0 |
| M-47 | 2 | 1 | 0 | 1 | 1 |
| M-63 | 1 | 0 | 1 | 0 | 0 |
| M-78 | 1 | 0 | 1 | 0 | 0 |

\**TnPhoA* mutants (Rahman et al 2002).

### Genome sequencing

A shotgun sequencing strategy was used for sequencing genomic DNA from the *P. alcalifaciens* 2939/90 and the four Tn*PhoA* mutants. Genomic DNA was extracted using the DNeasy blood and tissue extraction kit (Qiagen). Sequencing libraries were prepared using the Nextera DNA sample preparation kit (Illumina) and the sequence read data were produced on either the Illumina NextSeq (paired-end, 150 base reads) or MiSeq (paired-end, 300 base reads) instrument. Long-read shotgun sequence data for *P. alcalifaciens* 2939/90 genomic DNA was generated from a sequencing library prepared using the Rapid PCR Barcoding Kit and run on the Oxford Nanopore (ONT) MinION instrument. Sequence read data for the mutants are available at the National Center for Biotechnology Information (NCBI) in Bioproject, PRJNA1073245 and the assembled closed genome sequence for *P. alcalifaciens* 2939/90 is available in NCBI Bioproject, PRJNA929094. The genome sequence was annotated at NCBI using Prokaryotic Genome Annotation Pipeline (PGAP) (Parks et al., 2015).

### Genome assembly

A genome sequence was produced from the Illumina read data using Spades v3.9 (Bankevich et al. 2012) for each of the Tn*PhoA* mutants. A closed genome sequence for *P. alcalifaciens* 2939/90 was assembled using dragonflye (https://github.com/rpetit3/dragonflye) on ONT long read data for the assembly; Illumina read data were used for correcting the ONT assembly using a read mapping approach. In all cases, a preliminary purity check and confirmation of taxonomic classification was performed on read data sets using kraken2 (https://github.com/DerrickWood/kraken2) with the GTDB kraken2 database (generated by https://github.com/leylabmpi/Struo2 using https://gtdb.ecogenomic.org/).

### Tn*PhoA* insertion site

The Tn*PhoA* insertion site(s) for each of the Tn*PhoA* mutants was determined using the first and the last 30 bases of the sequence of Tn*PhoA* (NCBI Accession, U25548.1) to screen for reads containing: CCGTTCAGGACGCTACTTGTGTATAAGAGTCAG (Bases 7701 to 7733, top strand of U25548) and TCCAGGACGCTACTTGTGTATAAGAGTCAG (Bases 1 to 30, reverse strand of U25548). The identified reads were aligned, and a consensus sequence was determined for each insertion site in each isolate.

### Bioinformatics and data resources

Available *P. alcalifaciens* genome sequences from NCBI were obtained (downloaded on 1 February 2024). In total, 52 *P. alcalifaciens* genomes sequences were identified (including strain 2939/90). The list comprises genomes identified as *P. alcalifaciens* on NCBI or GTDB (https://gtdb.ecogenomic.org/) and excluded metagenome assembled genomes and poor-quality genomes (i.e., from mixed isolates or incomplete genomes). Summary information about these genomes is presented in Table S1. Type strain information was obtained from LPSN (https://lpsn.dsmz.de/) and a list of *P. alcalifaciens* genomes checked for genome completeness and taxonomic classification was obtained from GTDB. A comparison of isolate genomic sequences using a k-mer based approach was conducted using Mashtree (https://github.com/lskatz/mashtree). Genome wide average nucleotide identity (ANI) was determined using FastANI (Jain et al., 2018). Phylogenetic trees were visualized using Figtree (https://github.com/rambaut/figtree). Prediction of T3SS effector proteins was performed using EffectiveDB (Eichinger et al., 2016).

## Results

### Assembly and annotation of *P. alcalifaciens* 2939/90

Assembly using dragonflye (v1.0.13) with Oxford Nanopore long- and Illumina short-read data produced a closed circular chromosome sequence (4,087,862 bp) and four circular plasmid sequences (p2939_90_1: 127,696; p2939_90_2 84,555; p2939_90_3: 58,284; p2939_90_4: 7,475). The genome sequence was annotated by NCBI (accession: GCF_029962585.1) using PGAP (version 6.6) which identified 4,201 genes including 3,970 protein coding genes and 121 pseudogenes located on the chromosome and plasmids.

### Assembly and characterization of the genome sequences of Tn*PhoA* mutants

Illumina read data were used to assemble genome sequences for each of the four Tn*PhoA* mutants of *P. alcalifaciens* strain 2939/90. Sequence and assembly information is presented in Table S2. A survey of antimicrobial resistance genes showed the presence of additional resistance genes (*blaTEM-1, aph(6)-Ic, ble* and *aph(3’)-IIa*) in each of the Tn*PhoA* mutants all of which were not present in the parent strain 2939/90; this is consistent with the integration of Tn*PhoA* in the mutants. A core genome comparison of the parent strain and mutants showed the maximum distance between any isolate pair was two single nucleotide polymorphisms (SNPs), indicating a close genomic relationship between the mutants and *P. alcalifaciens* strain 2939/90.

### Tn*PhoA* insertions

The location of Tn*PhoA* insertions in the genome of the Tn*PhoA* mutants was determined by searching for reads that contained the sequence (last 50 bases, see Materials and Methods for sequences) at either end of the Tn*PhoA* element. The locations shown in Table 3 for each of the Tn*PhoA* mutants is with reference to the closed genome sequence for *P. alcalifaciens* 2939/90 (Accession: GCF_029962585.1).

**Table 3.**
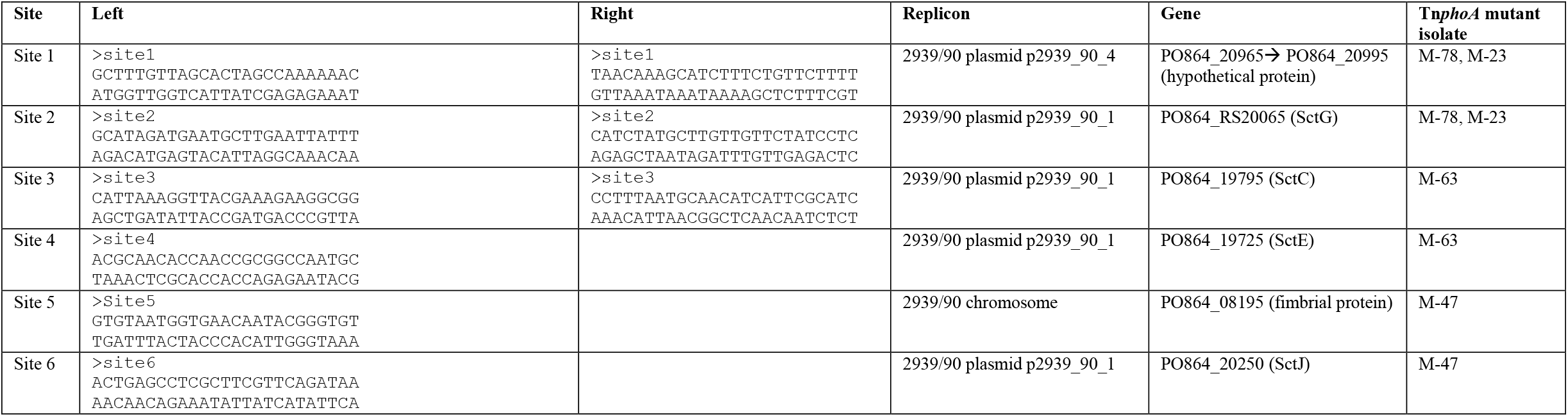
Tn*PhoA* insertion sites in mutant isolates of *P. alcalifaciens* 2939/90.

Each of the four Tn*PhoA* mutants carried two copies of the Tn*PhoA*. Mutants M-23 and M-78 were isogenic. Each of the four Tn*PhoA* mutants had at least one Tn*PhoA* inserted in plasmid p2939_90_1, with Tn*PhoA* mutant M-63 having both Tn*PhoA* copies in p2939_90_1. The chromosomal Tn*PhoA* insertion in Tn*PhoA* mutant M-47 interrupts a gene related to fimbriae biosynthesis, while for Tn*PhoA* mutants M-78 and M-23, the Tn*PhoA* insertion in plasmid p2939_90_4 interrupts a gene that may play a role in DNA conjugation. The insertion of Tn*phoA* in plasmid 1 of both M-78 and M-23 interrupts the gene encoding pilotin (SctG).

### Tn*PhoA* inserts in p2939_90_1 have a predicted role in type III secretion

Plasmid p2939_90_1 is 127,696 bp and is predicted to encode 115 proteins. A locus extending from bp 119,200 to bp 127,696 and then from bp 1 through to bp 22,805 contains genes encoding proteins that are predicted to be part of a type III secretion apparatus. All Tn*PhoA* mutants contain at least one Tn*PhoA* insertion site in this region of p2939_90_1 (see Table S3 for predicted gene location and function along with the genes which were interrupted by Tn*PhoA* in specific mutants).

### Two type III secretion apparatus loci in the *P. alcalifaciens* strain 2939/90 genome

Examination of the annotated chromosome of *P. alcalifaciens* strain 2939/90 identified another locus that is predicted to encode components of a type III secretion apparatus. The genes located in the respective type III secretion apparatus loci on the chromosome and on plasmid p2939_90_1 are shown in Table 4 (details of the location of the type III secretion apparatus genes on the chromosome are shown in Table S4).

**Table 4:**
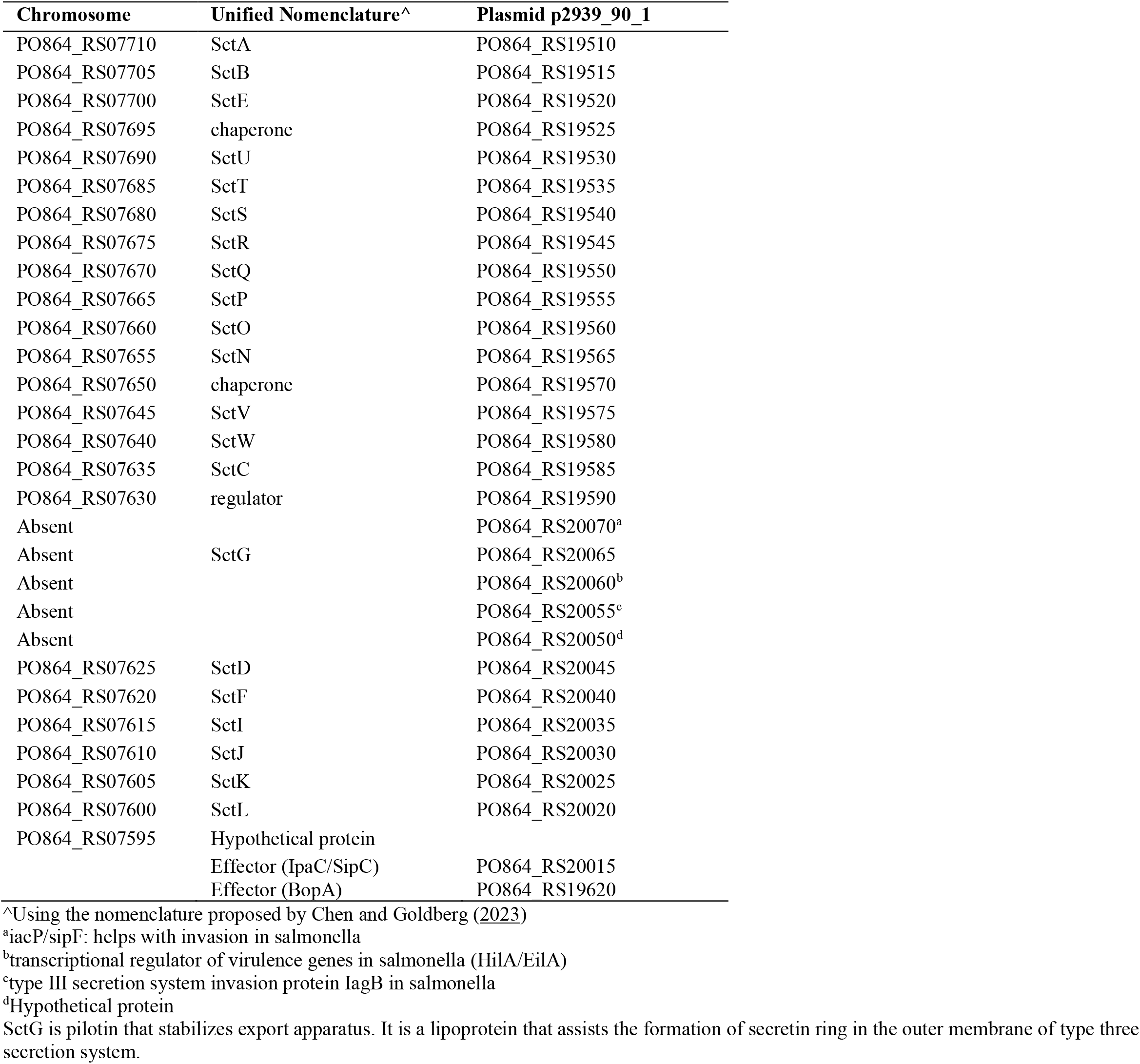
Genes encoding type III secretion apparatus proteins and some effectors in strain 2939/90.

### Distribution of the type III secretion apparatus loci in *P. alcalifaciens*

Taking the 52 available *P. alcalifaciens* genome sequences including the genome of strain 2939/90, we inferred genomic relationships among sequences using Mashtree (k-mer difference approach). A phylogenetic tree summarizing the inferred relationship is presented in Figure 1. We observed two major clades and identified these as Group A and Group B; strain 2939/90 is part of Group A. Using the region from 1,619,656 to 1,641,639 on the strain 2939/90 chromosome (type III secretion apparatus locus; see Table S4) as query, we determined using Blast that this locus was present in each of Group A genome sequences. At a lower average nucleotide sequence identity (∼85%), we detected a related locus in each of the Group B isolates; examination of the closed genome sequence for isolate 2019-04-29291-1-1 (Group B) showed a type III secretion apparatus locus with the same gene layout as for strain 2939/90 and with gene synteny in the regions flanking the locus between these genome sequences. The ANI between the Group A and Group B chromosomal genome sequence ranged between 88% and 89%; a similar rate of divergence between the Group A and Group B chromosomal type III secretion apparatus locus and the whole genome as well as genomic synteny suggested these chromosomally located type III secretion apparatus loci are orthologous and have been inherited vertically in *P. alcalifaciens*.

**Figure 1.**
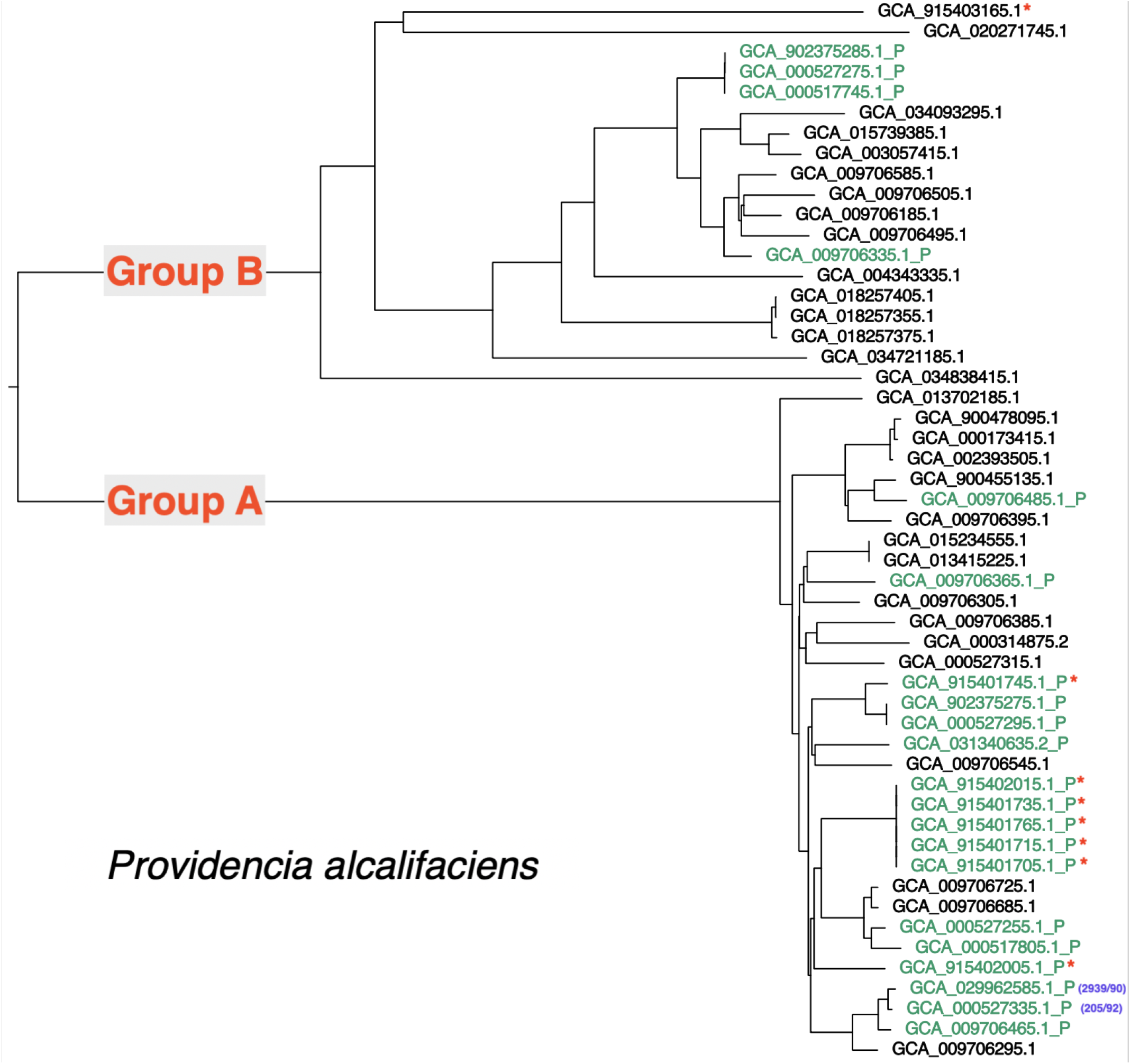
A phylogenetic tree showing the inferred relationship among the 52 available *P. alcalifaciens* genome sequences. The relationship was inferred using Mashtree. The tree shows two main clades of isolates (labelled Group A and Group B). Genome sequences were identified by the GenBank assembly accession numbers. Genome sequences containing the p2939_90_1 type three secretion system have a “_P” suffix and the taxon label is coloured green. Sequences that were part of the Norwegian *P. alcalifaciens* outbreak in dogs are identified with a red asterisk (Jorgensen et al., 2021). Strains 2939/90 and the related strain 205/92 (Albert et al., 1995) are identified with a purple text.

Again, using Blast and this time using the type III secretion apparatus locus located on plasmid p2939_90_1 as query, we investigated the distribution of this locus among the 52 available *P. alcalifaciens* genome sequences. Nucleotide Blast showed that there was no significant nucleotide similarity between the chromosomal and plasmid borne type III secretion apparatus locus; a broader search of the NCBI nr nucleotide database reveals this locus is present in sequence from *Providencia* and probably exclusively *P. alcalifaciens*. Constraining the e-value to e-100, the chromosomal type III secretion apparatus locus was not detected and a near identical locus was detected in a subset of the 52 genome sequences. Sequences containing the locus are shown in Figure 1. In total, 21 genome sequences contained the locus, with most being Group A genome sequences (17/21).

### Type III secreted effector protein prediction

A survey of type III secreted effector proteins encoded on the *P. alcalifaciens* strain 2939/90 was performed using EffectiveDB; a summary of the number of effector proteins predicted to be encoded on each replicon is shown in Table 5. A total of 26 predicted effectors were encoded on p2939_90_1. Among the effector proteins on this plasmid predicted using EffectiveDB (see Table S3), there are three effectors that would be predicted by protein similarity to a characterized effector protein (PO864_RS19515, related to the IpaC/SipC family of effector proteins; PO864_RS20070, related to IacP family of effector proteins and PO864_RS19620, related to BopA family of effector proteins), although, most predicted effector proteins were classified as hypothetical by protein sequence similarity. The distribution of these predicted effector genes among the 21 genomes carrying the p2939_90_1 T3SS locus is summarized in Table S5 and shown in detail in Table S6).

**Table 5.** Density of predicted T3SS effector CDS on each replicon of *P. alcalifaciens* strain 2939/90.

| Replicon | T3SS Effector (High) | T3SS Effector (Low) | Total CDS | Percent of CDS predicted to contain T3SS effectors |
| --- | --- | --- | --- | --- |
| Chromosome | 427 | 209 | 3580<sup>^</sup> | 17.8% |
| p2939_90_1 | 20 | 6 | 89<sup>^</sup> | 29.2% |
| p2939_90_2 | 5 | 3 | 73 | 11.0% |
| p2939_90_3 | 3 | 4 | 47 | 14.9% |
| p2939_90_4 | 0 | 0 | 8 | 0.0% |

<sup>^</sup>Genes encoding Type III secretion apparatus not included in the count.

## Discussion

Four Tn*PhoA* mutants of *P. alcalifaciens* strain 2939/90 were previously characterized and shown to have negligible invasion and actin condensation in HEp-2 cells (Rahman et al., 2002). Genomic characterization of these mutants has shown that these mutants have genomes that are highly related to the parental *P. alcalifaciens* 2939/90 strain (two or fewer core genome SNPs) and that the change in antimicrobial resistance genotype in the mutants is consistent with the insertion of Tn*PhoA* element into a replicon in each of the mutants. Interestingly, we determined that there were two Tn*PhoA* elements inserted in each of the mutants. While this is at odds with the observations made by Rahman et al (2002) where mutants were reported to contain a single insertion, additional insertion would have occurred during the subsequent propagation of the mutants.

Determination of the insertion sites showed that mutants M-78 and M-23 had identical Tn*PhoA* insertions (Table 3) and are therefore isogenic mutants. Each of the mutants had a Tn*PhoA* on p2939_90_1 except M-63 which had two Tn*PhoA* insertions on p2939_90_1. The remaining insertion sites were located on the chromosome or p2939_90_4. Based on the similarity of the cell culture phenotype seen in all four mutants, it could be assumed that the same function was being impacted by the Tn*PhoA* insertions in each of the mutants. Examination of the insertion points on p2939_90_1 and the predicted genes encoded at the points of insertion showed that all insertions on p2939_90_1 interrupted the genes involved in the type III secretion and all these genes were part of a type III secretion apparatus locus (Table 3).

A Type III secretion apparatus locus was also found on the chromosome of strain 2939/90; an orthologous locus was found on all 52 sequenced *P. alcalifaciens* genomes. This locus is likely to produce a type III secretion apparatus that is independent of the type III secretion apparatus encoded by the locus on p2939_90_1. This is supported by the observation that the p2939_90_1 locus was not present in all sequenced *P. alcalifaciens* genomes, and its distribution is not monophyletic (see Figure 1), potentially indicating horizontal movement of this locus. Both the chromosomal and plasmid p2939_90_1 loci contain all the required components to form a type III secretion apparatus (Chen and Goldberg, 2023), with the p2939_90_1 locus containing some additional, non-essential genes for the formation of a type III secretion apparatus. Moreover, we predict that as many as 26 effector proteins are encoded by p2939_90_1 which are likely to require the apparatus encoded on the p2939_90_1 for secretion, independent of the omnipresent chromosomally encoded type III secretion apparatus. By protein similarity, several proteins that are effectors in other pathogenic bacteria were detected in *P. alcalifaciens*. These included IagB, IacP/SinF, IpaC/SipC, BopA, EspG domain-containing protein, and TcdA/TcdB catalytic glycosyltransferase domain-containing protein. IagB and IacP/SinF are invasion proteins in salmonella (Miras et al., 1995; Kim et al., 2011), and IpaC/SipC is involved in invasion of epithelial cells by shigella/salmonella (Osiecki et al., 2001). Presence of these three invasive proteins in *P. alcalifaciens* strain 2939/90 is enough proof of their roles in invasive diarrhea. BopA is an effector protein secreted by *Burkholderia pseudomallei* via the type three secretion system, and it has been shown to play a crucial role in the escape of the bacterium from autophagy (Yu et al., 2016). EspG is an effector protein shared by enteropathogenic *Escherichia coli*, enterohaemorrhagic *E. coli* and shigella. It causes microtubule destabilization and cell detachment (Dean et al., 2010). TcdA and TcdB are primary virulence factors of *Clostridium difficile*. They enter and disrupt host cell function by glucosylating and thereby inactivating key signaling molecules within the host (Lacy and Alvin, 2017). Characterization of the function of the predicted effector proteins on p2939_90_1 seems to be a logical next step to investigate the complete diarrheagenic properties of *P. alcalifaciens*.

Thus, *P. alcalifaciens* 2939/90 carried two type III secretion systems (T3SSs). There are other pathogenic bacteria that are known to harbor more than one T3SS. These include *Salmonella enterica* (Hensel et al., 1995), *Yersinia enterocolitica* (Foultier et al., 2002), enterohaemorrhagic *Escherichia coli* O157:H7 (Hayashi et al., 2001), *Vibrio parahaemolyticus* (Park et al., 2004) and *Burkholderia pseudomallei* (Rainbow et al., 2002; Stevens et al., 2002). Both T3SSs were reported to be functional in *S. enterica* (Foultier et al., 2002) and *V. parahaemolyticus* (Park et al., 2004).

The invasive phenotype associated with strain 2939/90 in cell culture and the circumstances of the isolation of this strain from a fatal human case of diarrhea strongly suggest that *P. alcalifaciens* lineages carrying the p2939_90_1 type III secretion apparatus locus are likely to be able to cause severe diarrheal disease. While causation is difficult to demonstrate in humans, *Canis lupus familiaris* (dog) is a host that may well be useful for the demonstration of causation. We note that the *P. alcalifaciens* isolates characterized in an outbreak and associated with acute haemorrhagic diarrhea in dogs (Jorgensen et al., 2021) carried the p2939_90_1 type III secretion apparatus locus (shown in Figure 1).

While *P. alcalifaciens* is a well-recognized part of the normal flora of many animals including humans, it is not unprecedented for certain lineages within a commensal species to cause severe diarrheal disease; a case in point is diarrheagenic *E. coli*. Even though *E. coli* is commensal flora, at least five subgroups are recognized primary diarrheal pathogens (Kaper et al., 2004). Similarly, *P. alcalifaciens* strains that possess a p2939_90_1 type plasmid that carries a type III secretion apparatus locus are likely to be diarrheagenic. Such strains can be considered enteropathogenic as opposed to non-pathogenic normal flora strains.

## Conclusions

This work identified a T3SS encoded on p2939_90_1 (encoding both secretion apparatus and effector proteins) that may act independently of chromosomally encoded T3SS. The p2939_90_1 encoded T3SS contributes to the invasion phenotype observed in *P. alcalifaciens* strain 2939/90. The characterization of *P. alcalifaciens* strain 2939/90 and the observation that the presence of the p2939_90_1 T3SS in other isolates is associated with diarrheal disease is a significant step towards recognizing that a lineage or subgroup of *P. alcalifaciens* is a causative agent of diarrheal disease. This enteropathogenic lineage should be included in diarrheal disease investigations. The identification of genetic determinants that play a central role in disease causation in *P. alcalifaciens* makes it feasible to differentiate diarrhea causing lineage from normal flora lineages. A unique sequence that may be present in the pathogenic locus (p2939_90_1 T3SS locus) may be useful in a PCR assay to detect enteropathogenic strains of *P. alcalifaciens*.

